# A comparative analysis of current phasing and imputation software

**DOI:** 10.1101/2021.11.04.467340

**Authors:** Adriano De Marino, Abdallah Amr Mahmoud, Madhuchanda Bose, Karatuğ Ozan Bircan, Andrew Terpolovsky, Varuna Bamunusinghe, Umar Khan, Biljana Novković, Puya G. Yazdi

## Abstract

Whole-genome data has become significantly more accessible over the last two decades. This can largely be attributed to both reduced sequencing costs and imputation models which make it possible to obtain nearly whole-genome data from less expensive genotyping methods, such as microarray chips. Although there are many different approaches to imputation, the Hidden Markov Model remains the most widely used. In this study, we compared the latest versions of the most popular Hidden Markov Model based tools for phasing and imputation: Beagle 5.2, Eagle 2.4.1, Shapeit 4, Impute 5 and Minimac 4. We benchmarked them on three input datasets with three levels of chip density. We assessed each imputation software on the basis of accuracy, speed and memory usage, and showed how the choice of imputation accuracy metric can result in different interpretations. The highest average concordance rate was achieved by Beagle 5.2, followed by Impute 5 and Minimac 4, using a reference-based approach during phasing and the highest density chip. IQS and R^2^ metrics revealed that IMPUTE5 obtained better results for low frequency markers, while Beagle 5.2 remained more accurate for common markers (MAF>5%). Computational load as measured by run time was lower for Beagle 5.2 than Impute 5 and Minimac 4, while Minimac utilized the least memory of the imputation tools we compared. ShapeIT 4, used the least memory of the phasing tools examined, even with the highest density chip. Finally, we determined the combination of phasing software, imputation software, and reference panel, best suited for different situations and analysis needs and created an automated pipeline that provides a way for users to create customized chips designed to optimize their imputation results.

## Introduction

Genome wide association studies (GWAS) remain one of the most critical and powerful methods of identifying key genes and variants that play a role in many common human diseases (The Wellcome Trust Case Control Consortium 2007, Uffelmann et al. 2021). Identification of disease-associated variants in GWAS is dependent on successful tagging of millions of common variants in the human genome, and the ability to make inferences about genotypes of rare variants which are often not in linkage disequilibrium (LD) with common variants (The Wellcome Trust Case Control Consortium 2007, Uffelmann et al. 2021). Commercial single nucleotide polymorphism (SNP) genotyping arrays can contain up to 2.5 million markers, but none provide complete coverage of the human genome (Schurz et al. 2019). Despite the advances of the last two decades which have led to increasingly rapid and extensive genotyping, it is still prohibitively expensive to obtain whole genome sequencing (WGS) for the tens of thousands of individuals in GWAS (Peterson et al. 2017, Quick et al. 2020). Individual GWAS may also use distinct chips with different markers. To combine these GWAS for meta analysis, we require a method by which to identify genotypes at all markers utilized in each of these studies (Zaitlen and Eskin 2010). Thus, we continue to rely on imputation, the process of probabilistically estimating non-genotyped alleles for individuals in GWAS samples.

**Genotype imputation** is a method that infers the alleles of un-genotyped single-nucleotide polymorphisms (SNPs) based on the linkage disequilibrium (LD) with directly genotyped markers using a suitable reference population (Marchini and Howie 2010). It is predicated on the idea that seemingly unrelated individuals from the human population sampled at random can share short stretches of DNA within chromosomes derived from a shared ancestor (Scheet and Stephens 2006). Imputation can be used to improve SNP coverage and increase the statistical power of GWAS (Pei et al. 2010; Malhotra et al. 2014). Genotype imputation also facilitates fine mapping of causal variants, plays a key role in the meta-analyses of GWAS, and can be utilized in downstream applications of GWAS such as estimation of disease risk (Das, Abecasis, and Browning 2018). However, an important limitation of imputation is that only variants that were previously observed in a reference panel can be imputed (Das, Abecasis, and Browning 2018). Furthermore, rare variants are often poorly represented in reference panels making accurate imputation of rare and infrequent variants difficult. In addition, the choice of whether to pre-phase the data can impact imputation. Finally, imputation accuracy, sensitivity and computational efficiency are greatly affected by the choice of imputation software or tool (Das, Abecasis, and Browning 2018).

Over the last twenty years, multiple research groups have developed and published a number of phasing and imputation models, the majority of which are based on the Li and Stephens Hidden Markov Model (HMM) (Li and Stephens 2003). First described in 2003, it was applied to haplotype estimation methods, termed “**phasing**”, and used to handle large stretches of chromosome where individual haplotypes share contiguous, mosaic stretches with other haplotypes in the sample (Scheet and Stephens 2006, Das, Abecasis, and Browning 2018). Unlike previous coalescent approaches, it was computationally tractable, and methods based on the Li & Stephens HMM were soon shown to be more accurate and efficient than other methods (Lunter 2019, Scheet and Stephens 2006). Landmark and popular phasing algorithms are listed in Table 1, as a brief tabular history of the field. Currently, the most commonly used Li and Stephens HMM-based software’s are **BEAGLE**, **EAGLE**, and **SHAPEIT** for phasing, and **BEAGLE**, **IMPUTE** and **MINIMAC** for imputation.

**Table 1.** A brief history of phasing and imputation tools.

|  | Software | Published | Based on | Features | Complexity |
| --- | --- | --- | --- | --- | --- |
| Phasing | PHASE v 1.0 (Stephens, Smith, and Donnelly 2001) | 2001 | Coalescent approximation | | quadratic $O(n^2)$ |
|  | HAPI-UR (Williams et al. 2012) | 2012 | Li & Stephens HMM | used windows of sites instead of specific markers; led to higher accuracy |  |
| | Eagle 2 (Loh et al. 2016) | 2016 | Li & Stephens HMM | pBWT on a large reference panel condensed into a set of compact tree structures that losslessly model haplotype structure | linear $O(nm)$ |
| Phasing & Imputation | fastPHASE (Scheet and Stephens 2006) | 2006 | Li & Stephens HMM | | linear $O(n)$ |
| | Beagle v. 1.0 (Browning & Browning 2007) | 2007 | Li & Stephens HMM | uses bifurcating tree structure (aka haplotype-cluster model) | linear $O(n)$ |
| | Beagle v. 2.0, 3.0 (Browning & Browning 2009, 2013) | 2009 | Li & Stephens HMM | uses bifurcating tree structure (aka haplotype-cluster model) | linear $O(n)$ |
| | Beagle v. 4.0 (Browning & Browning 2018) | 2018 | Li & Stephens HMM | abandoned bifurcating model to adopt a flexible choice of haplotypes for reference similar to IMPUTE 2 | linear $O(n)$ |
|  | Beagle v. 5.2 (Browning & Browning 2021) | 2021 | Li & Stephens HMM | Introduction of progressive phasing algorithm to handle hundreds of millions of markers |  |
| | IMPUTE 2 (Howie, Donnelly and Marchini 2009) | 2009 | Li & Stephens HMM | flexible choice of haplotypes for reference panel; quadratic computational complexity meant inefficient | quadratic $O(n^2)$ |
| | IMPUTE 4 (Bycroft et al. 2018) | 2018 | Li & Stephens HMM | speed up haplotype imputation step | quadratic $O(n^2)$ |
| | IMPUTE 5 (Rubinacci, Delaneau and Marchini 2020) | 2019 | Li & Stephens HMM | uses positional BWT to choose haplotypes for each window | linear $O(nm)$ |
| | MACH (Li et al. 2010) | 2010 | Li & Stephens HMM | An iteratively updated phase of each study sample | quadratic $O(n^2)$ |
| | SHAPEIT 1 (Delaneau, Marchini and Zagury 2012) | 2011 | Li & Stephens HMM | flexible choice of the panel but computationally efficient | linear $O(n)$ |
| | SHAPEIT 2 (Delaneau and Marchini 2014) | 2013 | Li & Stephens HMM | combined best aspects of SHAPEIT 1 and IMPUTE 2 to increase accuracy and efficiency | linear $O(n)$ |
| | SHAPEIT 3 (Marchini et al. 2016) | 2016 | Li & Stephens HMM | increased scalability from SHAPEIT 2 | linear $O(n)$ |
| | SHAPEIT 4 (Delaneau et al. 2019) | 2018 | Li & Stephens HMM | pBWT to choose haplotypes for local window | linear $O(nm)$ |
| Imputation | Minimac (Howie et al 2012) | 2012 | Li & Stephens HMM | pre-phased imputation | linear $O(nm)$ |
| | Minimac 2 (Fuchsberger, Abecasis, and Hinds 2015) | 2014 | Li & Stephens HMM | | linear $O(nm)$ |
| | Minimac 3 (Das et al. 2016) | 2015 | Li & Stephens HMM | state-space reduction to reduce computational complexity and cost | linear $O(nm)$ |
| | Minimac4 (Júnior et al. 2021) | 2018 | Li & Stephens HMM | | linear $O(nm)$ |

A timeline and brief description of landmark and popular phasing and imputation algorithms and their computational complexities

Imputation accuracy is measured by several key sets of metrics which can be classified into two overarching types: statistics that compare imputed genotypes to ‘gold standard’ genotyped data and statistics produced without reference to true genotypes (Ramnarine et al. 2015). Concordance rate, squared correlation R^2^, and Imputation Quality Score (IQS) are examples of the first type (Candelaria Vergara 2018, Ramnarine et al. 2015). In practice, the purpose of imputation is to predict SNPs for which we do not have genotyped data; statistics of the second type are typically relied upon during imputation, and generally output by the various imputation programs. Although the rapid increase in the number of deeply sequenced individuals will soon make it possible to assemble increasingly large reference panels that greatly increase the number of imputable variants, the choice of phasing and imputation software currently has a significant impact on accuracy (Herzig et al. 2018). While several studies have evaluated and compared imputation models, or phasing models, or imputation models in combination with different reference panels, no recent studies have compared imputation and phasing algorithms in combination with different reference panels, in tandem, and evaluated the relative computational efficiency and accuracy of each combination (Sariya et al. 2019, Herzig et al. 2018).

In this study, we evaluate the latest versions of the most commonly used tools for phasing and imputation in terms of accuracy, computational speed and memory usage, using 2 different versions of the 1kG project as reference panels and three different microarray chip datasets as inputs. We combine each tool for phasing with a method for imputation to understand which combination achieves the best overall results and which method is the best at imputing rare variants. Our goal was to determine the combination of phasing and imputation software and reference panel that is best suited for different situations and needs.

## Methods

### Chip Data

We used three different chip datasets with differing marker density and input dataset sizes. The first chip dataset (**Affymetrix**) was composed of 3450 unrelated individuals from The 1000 Genomes Project genotyped with the Affymetrix 6.0 900K array (Affymetrix, ThermoFisher), the second (**Omni**) of 2318 unrelated individuals from the 1000 Genomes Project genotyped with the Omni 2.5 chip by Illumina 2.4 Million unphased SNP markers, and the third one (**Customized**) was a subset of the first two chips and consisted of the intersection of the first two chips with another chip, GSA version 3 with direct-to-consumer booster by Illumina (Fig. 1). This Customized chip is the intersection of commonly used chips, resulting in a low-density chip with fewer overall sites, to allow us to assess imputation and phasing accuracy when the input data is limited to a relatively small number of SNPs.

**Fig. 1.**
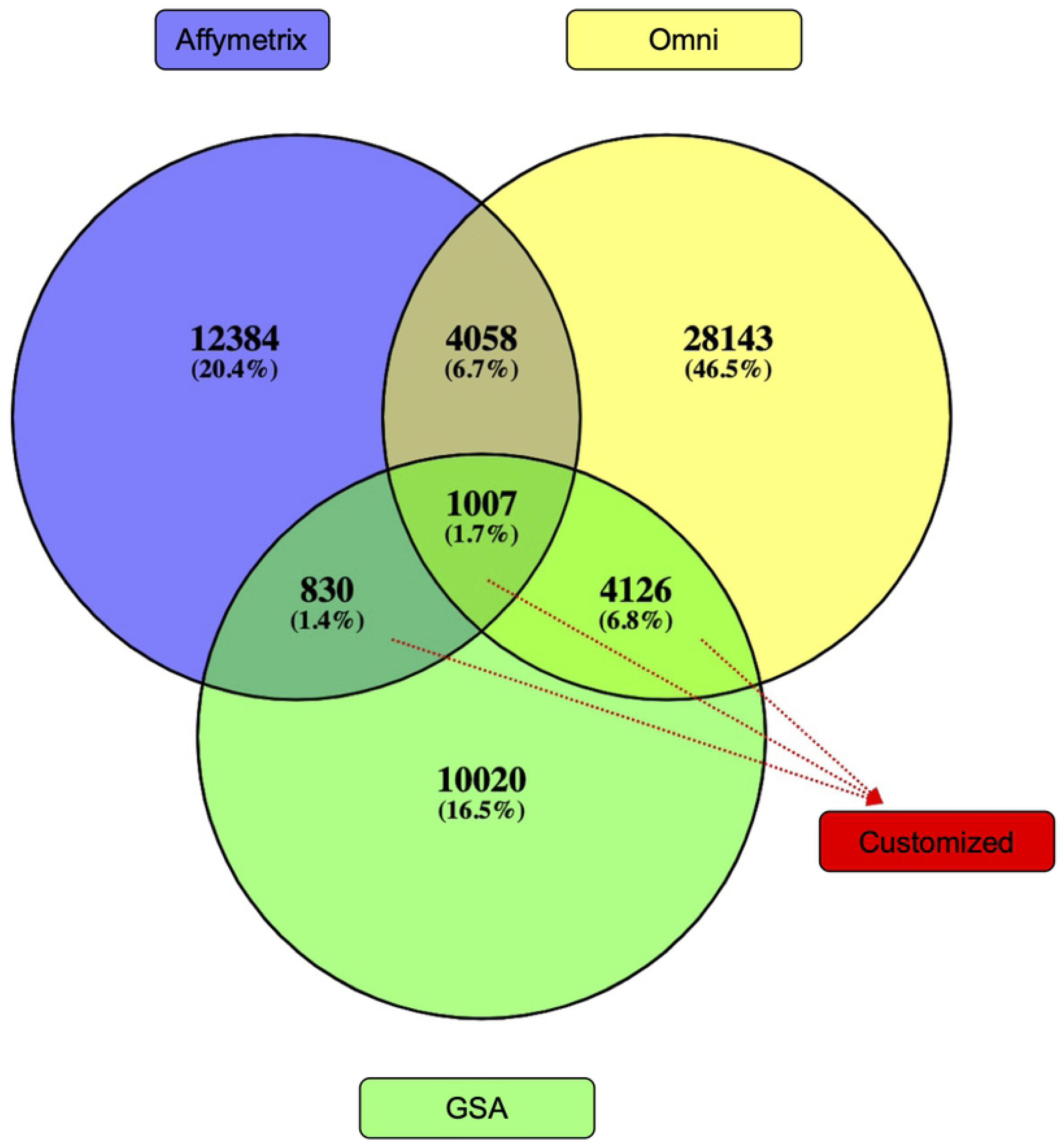
Chip data used to assess imputation and phasing accuracy and origin of the customized chip. Affymetrix, Omni and Customized chips. SNP numbers for chromosome 20 are shown. Customized chip data was obtained from the intersection of the first two chips with the Eurofins chip.

Fig. 2 describes the preparation of chip datasets for analysis. Data from Affymetrix and Omni chips were normalized using BCFtools (Petr Danecek 2021). Chip data was processed separately for each chromosome, which was renamed numerically with the ‘chr’ tag to match the reference panel. Chromosome 20 was chosen for use in all downstream analyses as it is generally representative of autosomal chromosomes. Sample data was converted to GRCh38 with Picard liftover (Picard Toolkit 2019), to match the assembly of the reference panels. We split multiallelic sites to record them as biallelic, left-normalized the variants to the reference genome, and removed duplicate variants. Finally, because Beagle does not allow skipping imputation of sporadic missing data, variants with missing genotype information were removed from the chip datasets and the WGS reference panels.

**Fig. 2.**
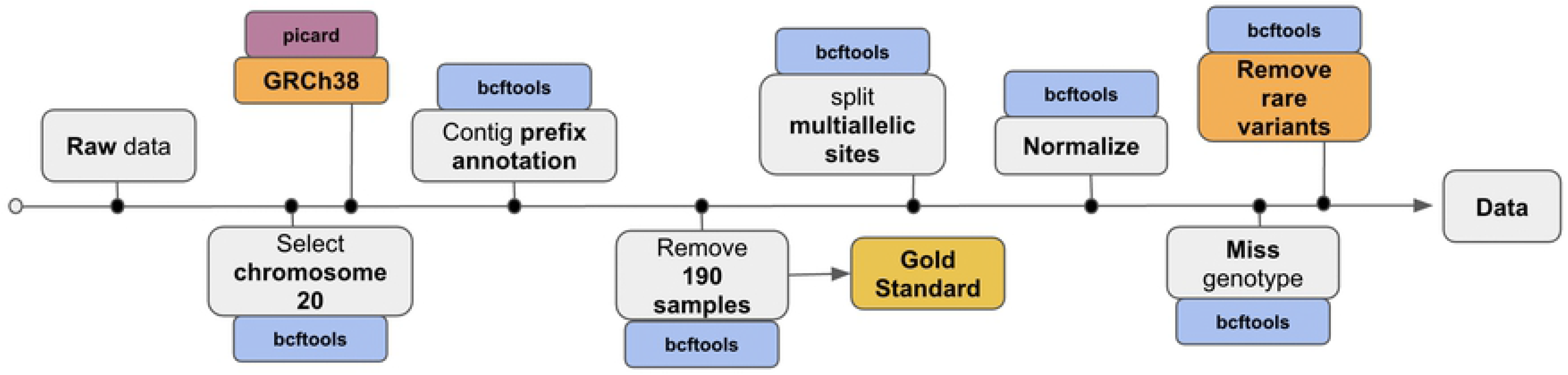
Pre-processing of the HD genotype chips and reference panels. Pre-processing of the HD genotype chips and reference panels downloaded from the International Genome Sample Resource (IGSR). Steps highlighted in orange are specific to the 1000GPphase3 reference panel only; all other steps were performed for both reference panels.

### Reference Panel Collection and Sample Selection

We drew our reference panels for imputation and phasing from the The 1000 Genomes Project (1000GP). We used the Phase 3 low coverage WGS which has a mean depth of 7X as one reference panel and the high coverage WGS, with a mean depth of 30x, as a second reference panel (1000 Genomes Project Consortium et al. 2010, 467; 2010, 491; 2015, 526; Sudmant et al. 2015). We refer to these as the 1000GP-Phase3 and 1000GP-30x reference panels.

We randomly selected 190 unrelated individuals taken from the set of 1686 individuals found in all three collections -- the Omni, Affymetrix and WGS 1000 Genomes Project sample collections (Sudmant et al. 2015) as shown in Fig 3. Our sample consisted of 5 males and 5 females per population, for 19 different populations and 5 super-populations (Fig 4.). These 190 individuals, and their relatives, were removed from the reference panels and used to create chip datasets for testing. Imputation accuracy was assessed by looking at the concordance between the imputed chips’ data and the whole genome sequences for these 190 samples.

**Fig. 3.**
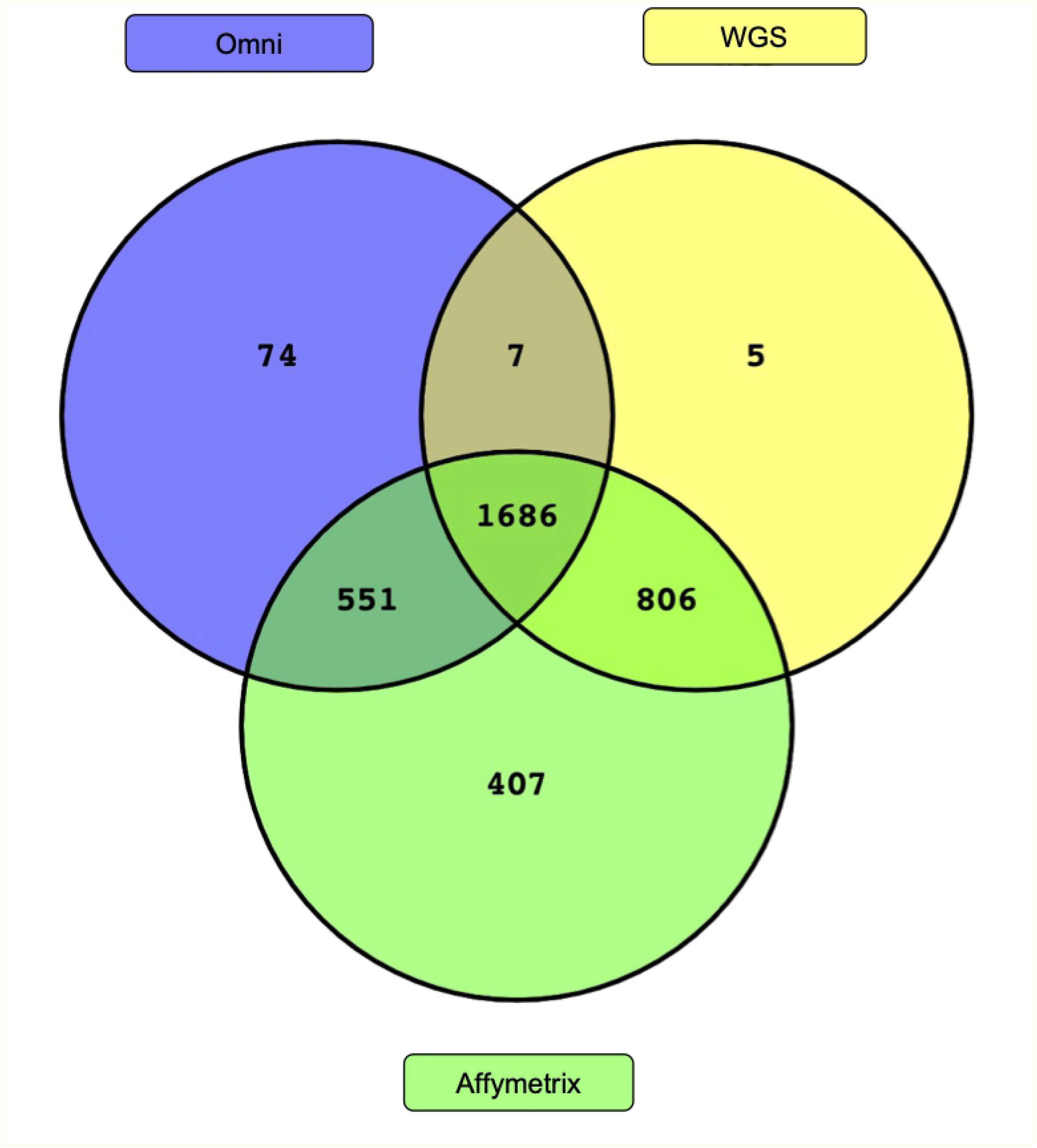
Shared individuals between HD genotype chips and reference panels. Individuals in common between the WGS Reference panels, Omni and Affymetrix chips.

**Fig. 4.**
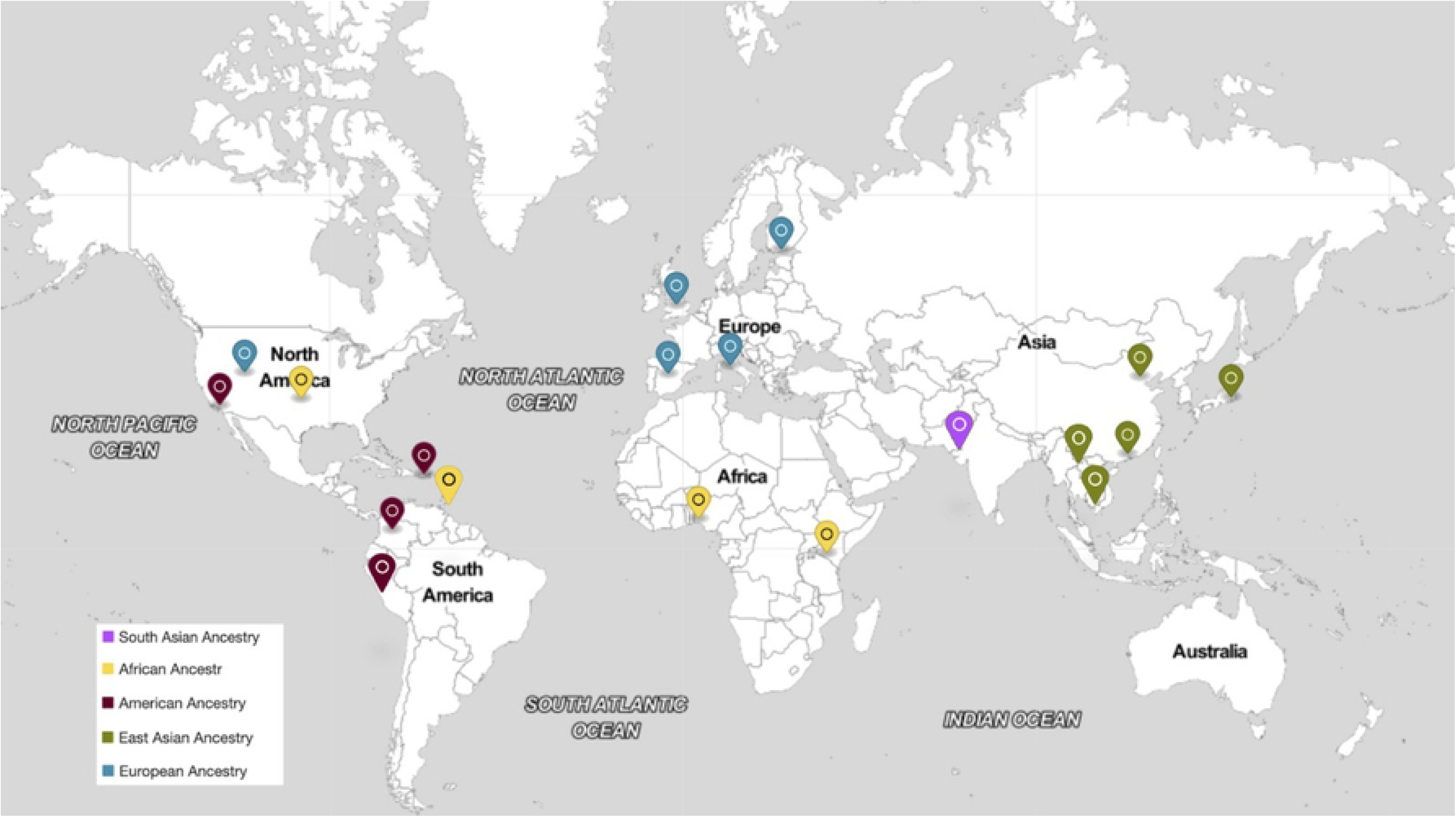
Origin of the target samples. Sample of 190 individuals belonging to 19 populations from 5 super populations selected for this study.

### Quality Control of Reference Panels

For both reference panels, we used BCFtools (Petr Danecek 2021) to split multiallelic sites, remove duplicates and missing data, and align variants to the reference genome. Both the 1000GP30x and 1000GPphase3 panels were preprocessed by prepending the contig name with the prefix ‘chr’. Two additional steps were performed for the 1000GPphase3 panel to convert it to GRCh38 with Picard liftover (Picard Toolkit 2019), and discard rare variant singletons and doubletons to evaluate if their removal increased imputation accuracy for common variants (MAF>5%). The workflow for the quality control and pre-processing of the reference panels is shown in Fig. 2.

### Phasing and Imputation Pipeline

The **Affymetrix**, **Omni** and **Customized** chips were used as inputs for 9 combinations of phasing and imputation tools to assess which combination performed best for our sample set (Fig. 5), using one of the two reference panels. Phasing was performed using both **reference-free** and **reference-based** approaches for each method, to compare their respective resultant imputation accuracy. This yielded a total of 108 combinations of input chip dataset, phasing tool, reference-based or reference-free phasing, imputation reference panel, and imputation tool (Supplementary Table 1).

**Fig. 5.**
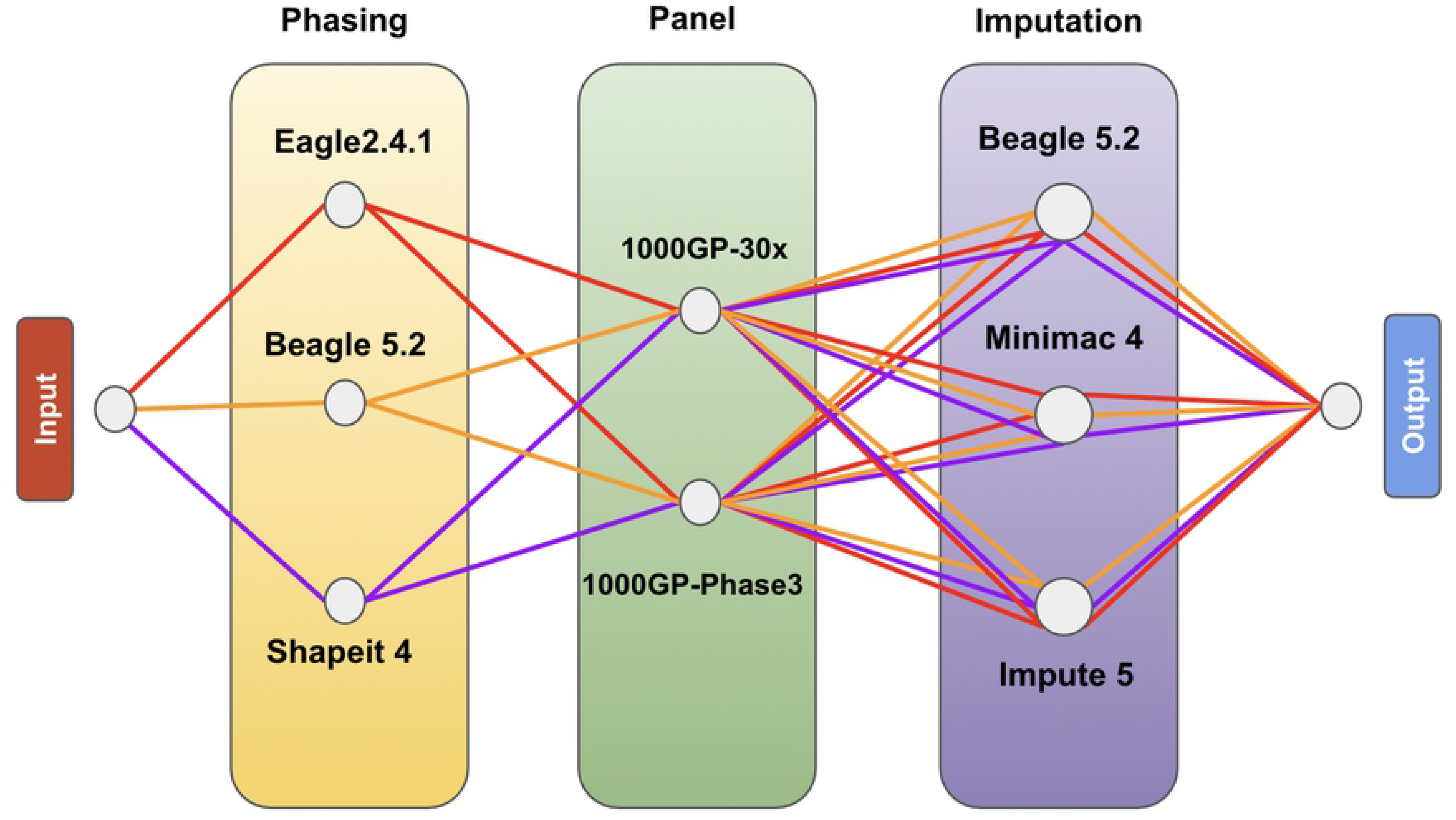
Workflow of the analysis, combinations tested. Each input chip dataset was analysed using the 36 combinations of 3 different phasing softwares, 2 phasing approaches, 3 imputation softwares, and 2 imputation reference panels.

The haplotype phasing softwares we compared are: **Eagle2 v2.4.1** (Loh et al. 2016), **Beagle5 v5.2** (Browning, Zhou, and Browning 2018), and **Shapeit4 v4.2.1** (Delaneau et al. 2019). All phasing software was launched with default parameters using 4 cores for each analysis on an Intel Corporation 82371AB/EB/MB PIIX4 ACPI 64-bit 32Gb RAM and the saved log file was used to evaluate the total run time. The imputation methods we tested are: **Beagle5 v5.2** (Browning, Zhou, and Browning 2018), **Impute5 v1.1.5** (Rubinacci, Delaneau, and Marchini 2020) and **Minimac4 v1.0.0** (Das et al. 2016).

Each input chip dataset was processed using **selphi.sh** (SELfdecode PHasing and Imputation) an automated pipeline built in bash that combines the phasing and imputation software and evaluates accuracy at each step to speed up the process of analysis and comparison. The inputs to the pipeline are the chip data file, a reference panel, the number of threads to use and the chromosome to process. The pipeline first checks that the correct version of the reference panel already exists for each imputation software to use and if the input file is available both in BCF format and in VCF format. This means that the original reference panel is converted to **bref3** for Imputation with Beagle5.2 using bref3.29May21.d6d.jar, to **m3mv** for Minimac4 using Minimac3 and to **imp5** for Impute5 using imp5Converter_1.1.5_static. If any of these files don’t exist, they are automatically created by the pipeline. After this initial check, the pipeline begins phasing the haplotypes using Eagle2.4.1, Beagle5.2 and Shapit4. Each of these softwares was run twice with default parameters, once with the reference and once without, using 4 threads on chromosome 20 with recombination rates drawn from the genetic map. This step generated 2 phased VCF files for each software, yielding a total of 6 phased VCF files. After phasing, VCF files were moved to imputation with Beagle5.2, Minimac4 and Impute5. All were run using default parameters with a genetic map for the recombination rate and 4 threads. There are options to speed up both Minimac4 and Impute5 but these tend to reduce the accuracy rate. To maximize the accuracy of each tool and preserve the validity of the comparison, we ran them with the default parameters, avoiding the steps required to optimize for computational load.

### Accuracy Measurement

Accuracy was assessed by comparing the imputation data resulting from each of the different combinations of phasing tool, imputation tool, and choice of reference, against the WGS dataset of the chosen 190 target samples. Variables considered were population/ancestry, sex, choice of tools, choice of reference, use of a reference panel, chip density, and the effect of MAF. We also looked at computational efficiency and memory usage. To check the effects of MAF on imputation accuracy, we used r^2^ as the metric of choice as it can distinguish between different MAF stratifications and is the most widely used metric for assessing imputation accuracy (Liu et al. 2013).

Accuracy was evaluated using a custom, faster version of the imputation accuracy calculation software available on github (Chen et al. 2020) that summarizes the accuracy metrics described in the work of Ramnarine et al. 2015 (Ramnarine et al. 2015). A detailed report with the concordance ratio (Po), F-measure score, square correlation (R^2^) and imputation quality score (IQS) was generated and written to the output file. To accurately assess IQS and R^2^ results, we removed all variants with MAF equal to 0 in our target population (allele count equal to 0) of 190 individuals from the analysis; IQS is zero when MAF is equal to zero, and is not indicative of accuracy or imputation quality. The entire code for accuracy metrics can be found in the script simpy.py (see section Data Available).

## Results

### Genotyping Data

After performing quality control on chromosome 20, 18,279 variants with a genotyping call rate of 100% remained in the Affymetrix chip dataset, and 37,334 variants with a genotyping call rate of 100% remained in the Omni Illumina dataset. In total, 5065 SNP markers overlapped between the two chips. The customized chip had 5913 markers shared between the Eurofins and the Affymetrix and Omni chips. The number of variants shared between the chip datasets and the 1000GP-30x panel (WGS) is shown in Fig. 6.

**Fig. 6.**
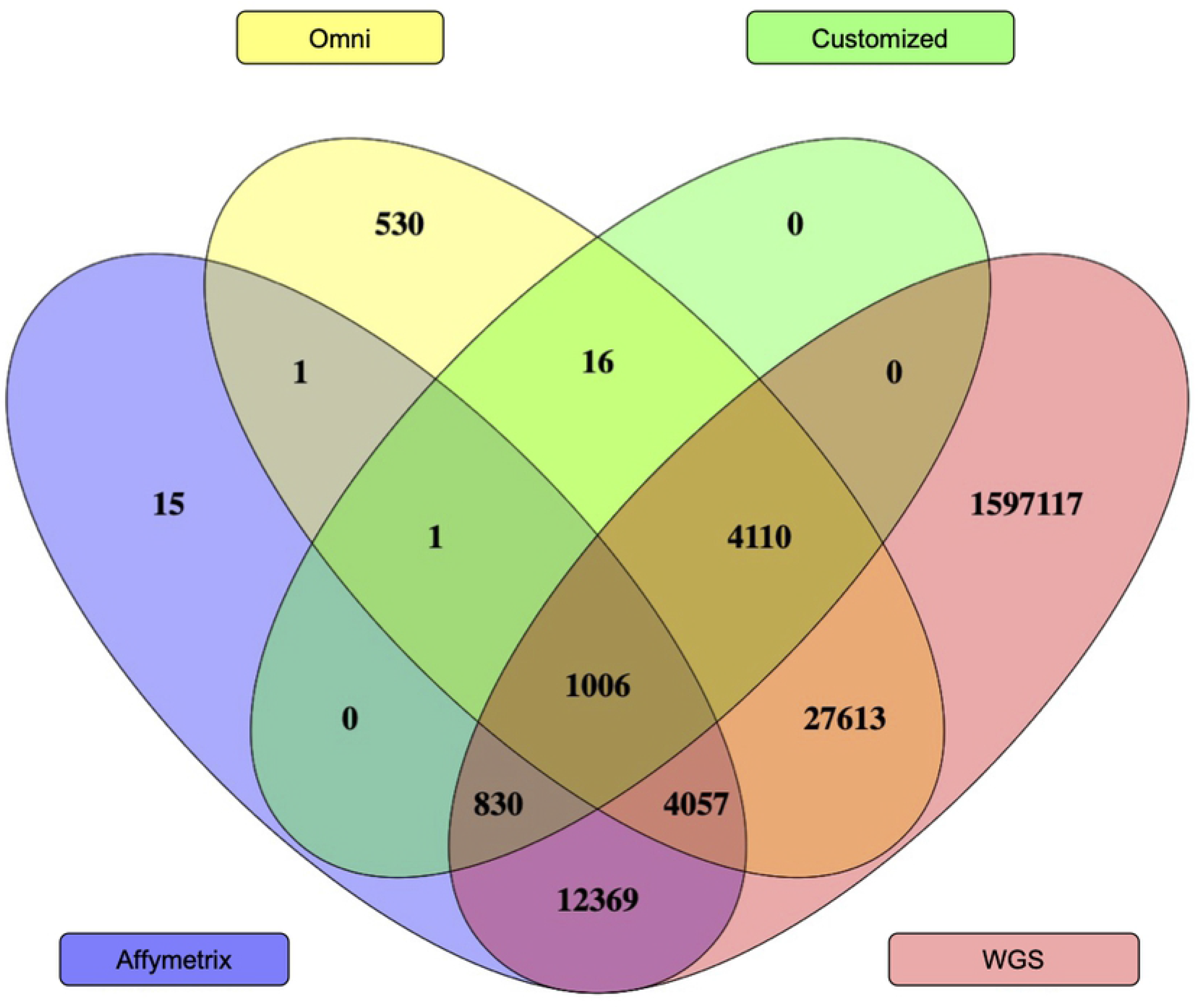
Number of shared variants between datasets. Variants on chromosome 20 shared between chips and the 1000GP-30x WGS reference panel.

### Imputation Accuracy

#### Minor Allele Frequency (MAF) And Reference Panel

We stratified variants based on MAF and assessed imputation accuracy for common, infrequent, and rare variants to obtain a more nuanced understanding of how well each combination of phasing-imputation tools performed (Table 2).

**Table 2.**
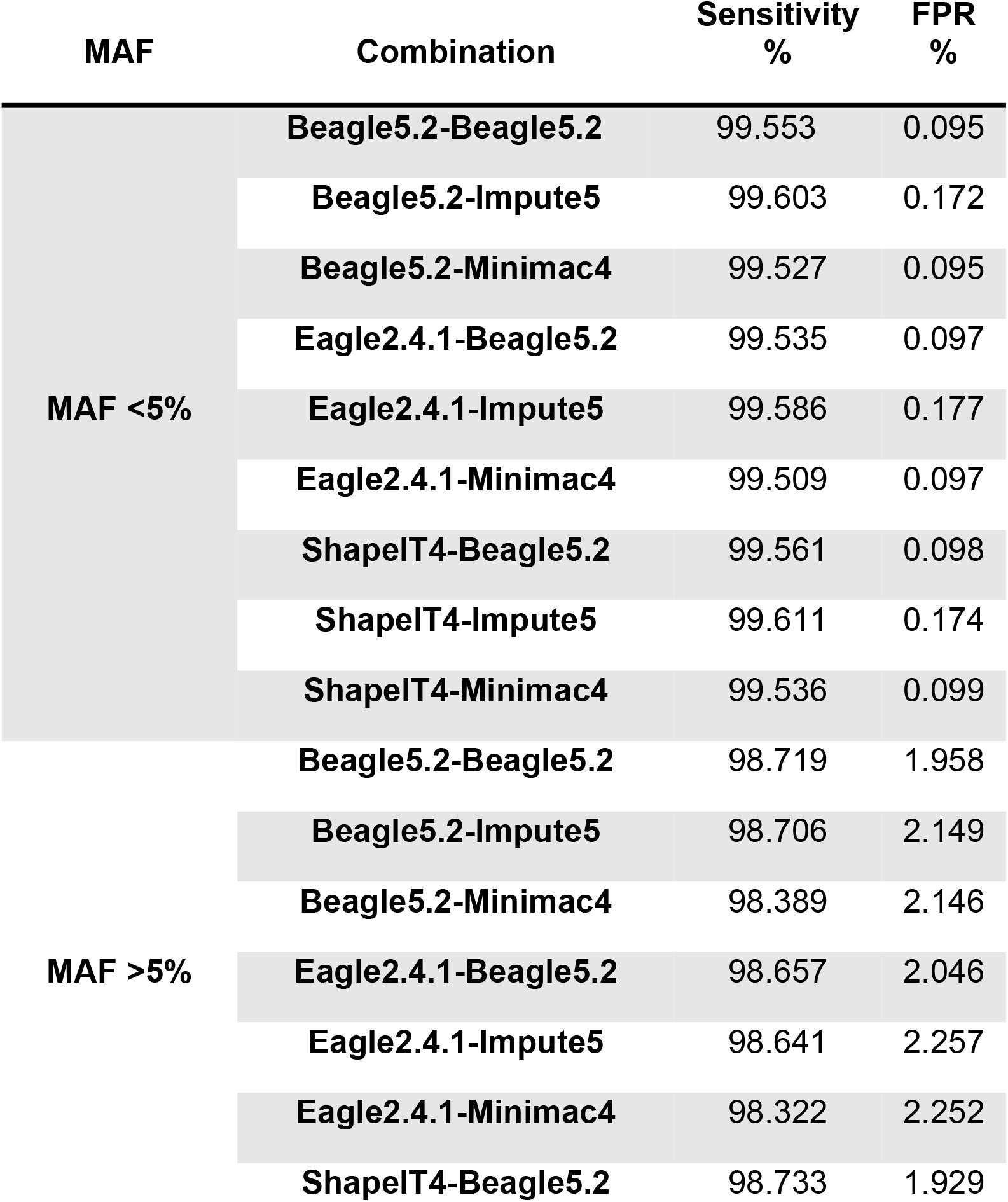
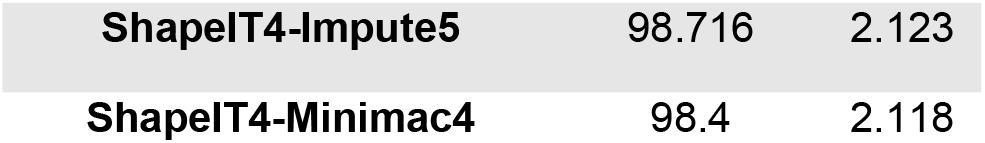
MAF-stratified comparison of phasing-imputation combinations.

| MAF | Combination | Sensitivity<br>% | FPR<br>% |
| --- | --- | --- | --- |
| MAF <5% | Beagle5.2-Beagle5.2 | 99.553 | 0.095 |
|  | Beagle5.2-Impute5 | 99.603 | 0.172 |
|  | Beagle5.2-Minimac4 | 99.527 | 0.095 |
|  | Eagle2.4.1-Beagle5.2 | 99.535 | 0.097 |
|  | Eagle2.4.1-Impute5 | 99.586 | 0.177 |
|  | Eagle2.4.1-Minimac4 | 99.509 | 0.097 |
|  | ShapIT4-Beagle5.2 | 99.561 | 0.098 |
|  | ShapIT4-Impute5 | 99.611 | 0.174 |
|  | ShapIT4-Minimac4 | 99.536 | 0.099 |
| MAF >5% | Beagle5.2-Beagle5.2 | 98.719 | 1.958 |
|  | Beagle5.2-Impute5 | 98.706 | 2.149 |
|  | Beagle5.2-Minimac4 | 98.389 | 2.146 |
|  | Eagle2.4.1-Beagle5.2 | 98.657 | 2.046 |
|  | Eagle2.4.1-Impute5 | 98.641 | 2.257 |
|  | Eagle2.4.1-Minimac4 | 98.322 | 2.252 |
|  | ShapIT4-Beagle5.2 | 98.733 | 1.929 |

|  |  |  |
| --- | --- | --- |
| ShapeIT4-Impute5 | 98.716 | 2.123 |
| ShapeIT4-Minimac4 | 98.4 | 2.118 |

A comparison of the sensitivity and false positive rate (FPR) of the imputation results, for each phasing-imputation combination, as stratified by MAF.

Based on the accuracy metric, the False Positive Rate (FPR), and the sensitivity, Beagle5.2 outperformed other phasing tools when MAF was greater than 5%, with ShapeIT4 a close second. However, for uncommon variants (MAF≤5%), ShapeIT4 was the better phasing tool, irrespective of imputation tool choice. For the imputation of uncommon variants (MAF ≤ 5%), Impute5 outperformed Beagle5.2 and Minimac, for each phasing tool combination. However, for common variants (MAF ≥ 5%), Beagle5.2 was superior. Similar results were obtained using r^2^ as the metric (Fig. 7). The best combination overall was ShapeIT4-Beagle5.2 imputing from the Omni chip dataset, with a reference-based phasing approach and imputing using the 1000GP-Phase3 reference panel, resulting in an average imputation r^2^ of 0.839 (S1 Table 1). On the other hand, for the 1000GP-30x reference panel, the best phasing and imputation tool combination was ShapeIT4-Impute5 using an Omni chip with reference-based phasing, resulting in an average imputation r^2^ of 0.728 (S1 Table 1).

**Fig. 7.**
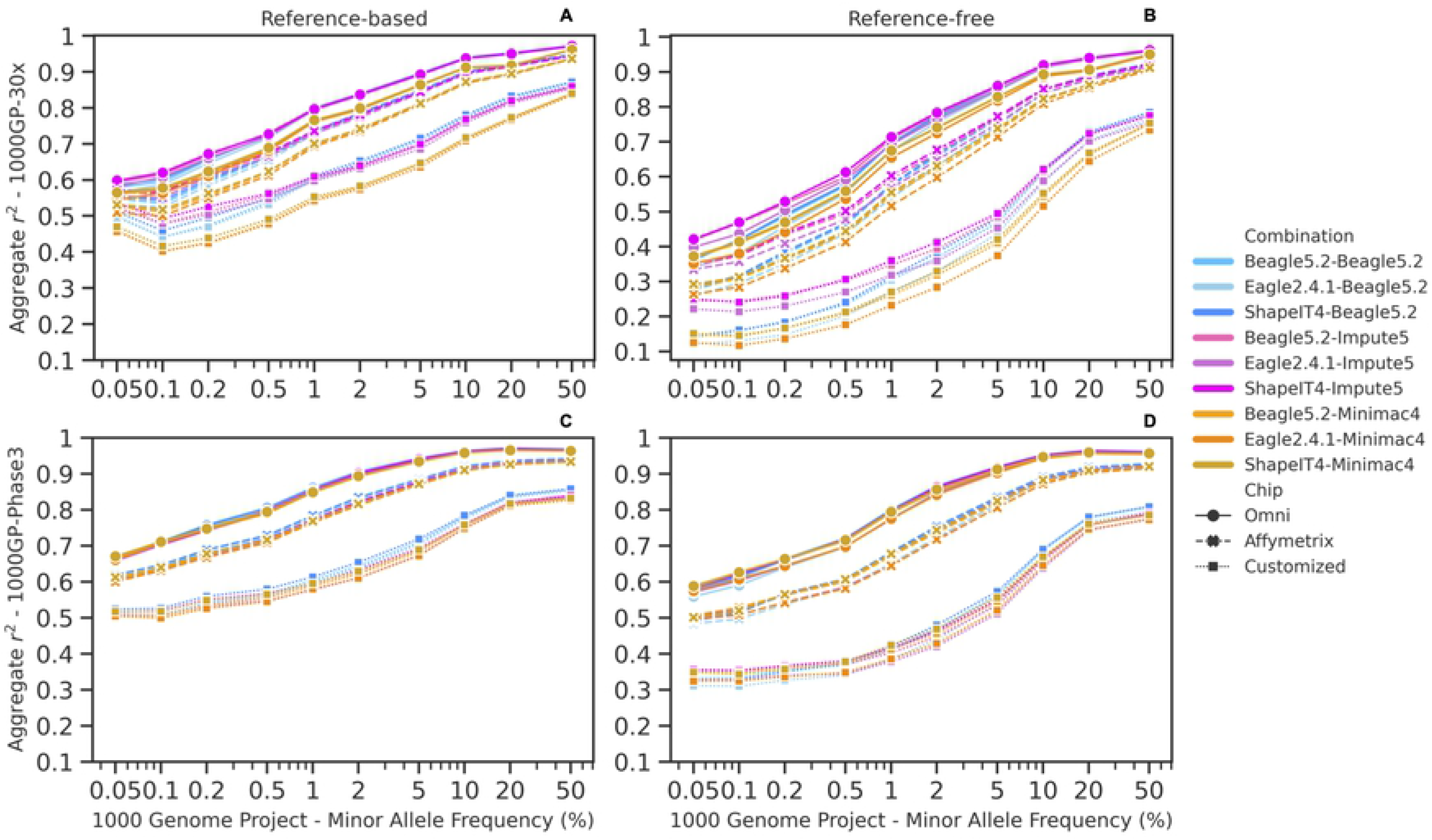
Imputation performance for chromosome 20 using 190 mixed population individuals with 2 reference panels and 2 phasing approaches. Blue colors indicate Beagle5.2, violets indicate Impute5 and oranges indicate Minimac4. The different input chip datasets are notated using the shape of the line: dashed for Affymetrix, continuous for Omni and dotted for the customized chip. (A) reference-based - 1000GP-30x, (B) reference-free - 1000GP-30x, (C) reference-based - 1000GP-Phase3, (D) reference-free - 1000GP-Phase3.

Fig. 8 depicts an increase in IQS with increasing MAF. Impute5 produced better results at lower MAF than either Beagle5.2 or Minimac4, while Beagle5.2 imputed better above 5% allele frequency. Ultra-rare variants were imputed badly with all available software.

**Fig. 8.**
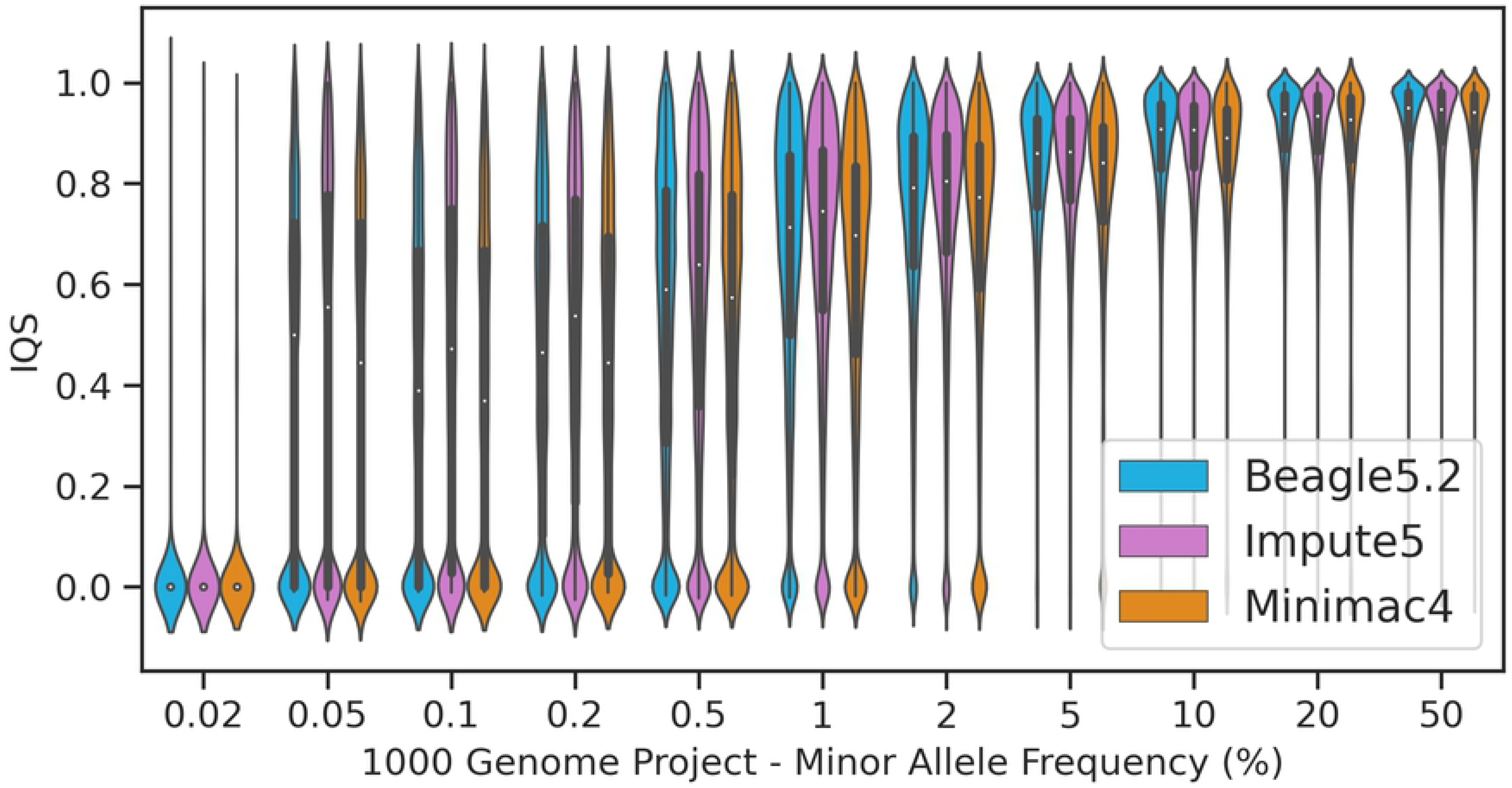
Evaluation of rare variants imputation. Violin plot. IQS is plotted against Minor allele frequency (MAF).

Choosing ShapeIT4 as the phasing tool for reference-based phasing, followed by any choice of imputation tool, resulted in the highest r^2^ for either imputation reference panel (S1 Table 1). For the Affymetrix and customized chips, ShapeIT4 remained the best choice of phasing tool for reference-free phasing, with respect to r^2^; for Omni, Beagle was the superior phasing tool. However, when we instead considered IQS as the metric of choice, both Beagle and ShapeIT4 performed equally well for reference-based phasing for higher density input chip datasets, but ShapeIT4 outperformed Beagle for the customized chip dataset, which had low chip density. For reference-free phasing, with respect to IQS, there was no clear winner between ShapeIT4 and Beagle (S1 Table 1).

To get a better overall representation of how MAF affects imputation accuracy and error rates, we plotted IQS against Error rate (Fig. 9), where each dot represents an imputed variant. The markers cluster according to their MAF and follow a waterfall trend. The results of this analysis are shown in Figure 9, which illustrates that IQS is generally higher and error rates overall lower for more common variants. Rare variants, with MAF<1%, tend to have lower IQS and higher error rates.

**Fig. 9.**
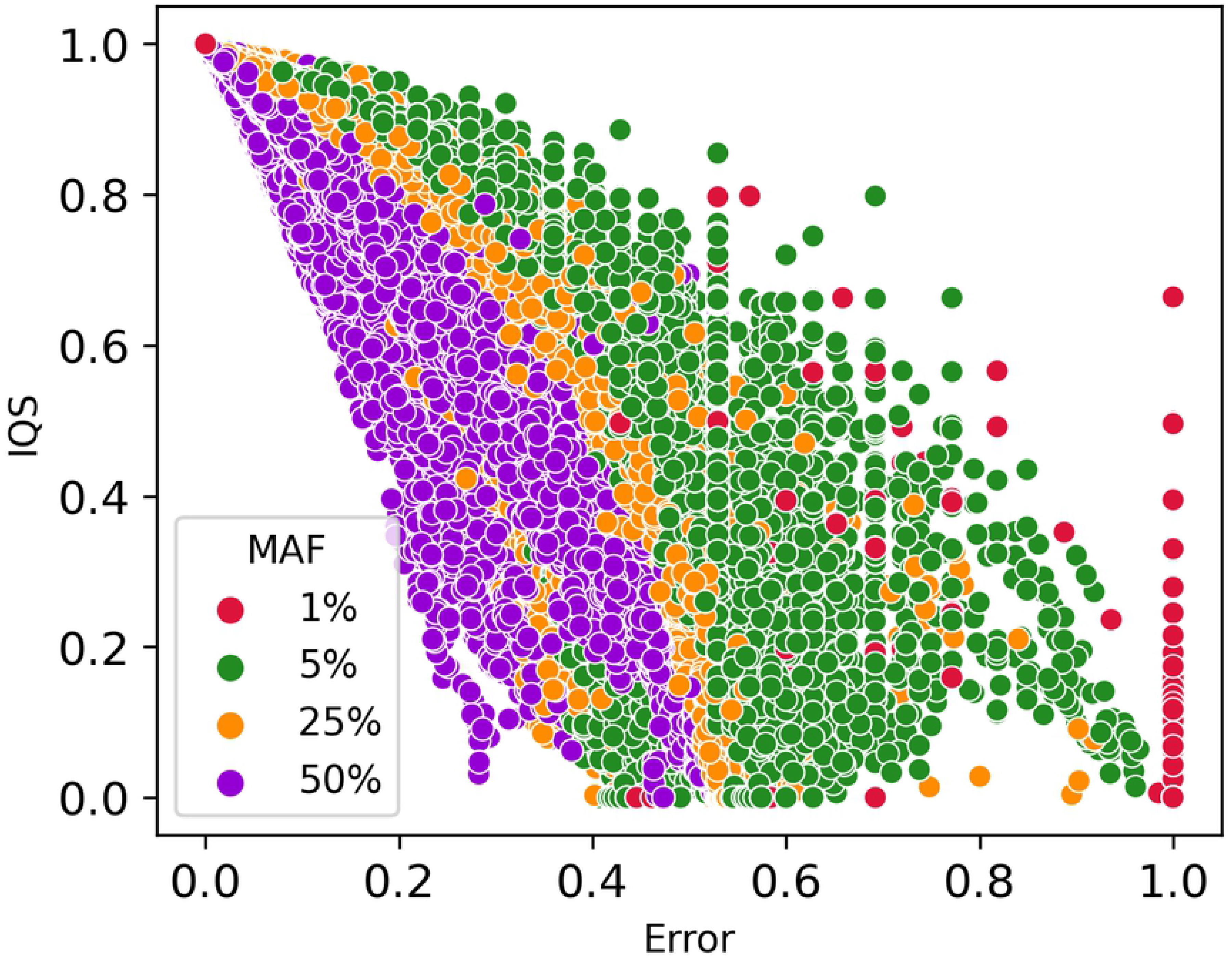
Minor allele frequency (MAF) Stratification of imputed variants. Dots are clustered following minor allele frequency stratification. The dots clustered in the right-down corner of the figure have low IQS and high Error rate, while dots in the left-high corner have high IQS and low Error rate. Each dot represents the average IQS and error rate for a specific marker imputed with one phasing tool-imputation tool combination.

#### Population, Sex, Chip Density, and Phasing Approach

Accuracy as measured by concordance (Po) was lowest in individuals of African ancestry, and highest in individuals of European and American populations--groups which both have significant recent European ancestry (Table 3). Furthermore, despite reaching similar average imputation accuracy, a greater proportion of EUR individuals had very high imputation accuracy compared with a progressively smaller proportion of target individuals with higher concordance for East Asian, American, African and South Asian ancestry, respectively (Fig. 10B). Thus, although we were able to reach similar mean imputation concordance for each of the different populations, imputation tools performed the best when applied to EUR populations and the worst for AFR and South Asian populations.

**Fig. 10.**
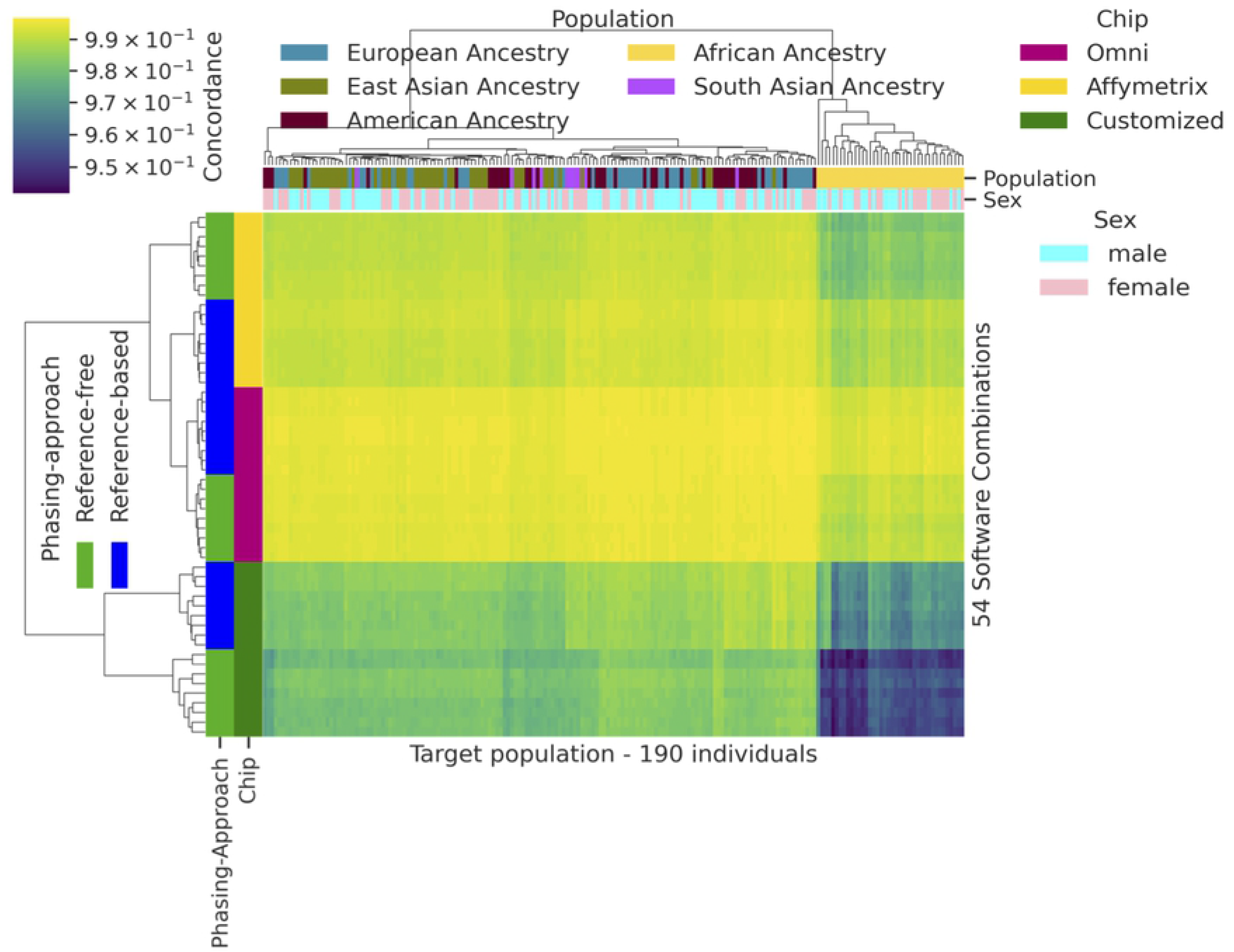
Imputation concordance rate over four different features. Stacked density plot of accuracy stratified by (A) sex; (B) superpopulation; (C) chip data; (D) phasing type (reference-free and reference-based).

**Table 3.** Accuracy for different Superpopulations.

| Superpopulation name | Mean | Std |
| --- | --- | --- |
| African | 0.984396 | 0.012613 |
| American | 0.993112 | 0.005104 |
| East Asian | 0.991575 | 0.004868 |
| European | 0.99274 | 0.004655 |
| South Asian | 0.991464 | 0.004989 |

Accuracy as measured by concordance (Po) of the imputation results for each of the five main superpopulations

Differences in imputation accuracy by population and phasing approach are shown in Figure 10. The reference-based approach produced better results than the reference-free approach, for most combinations of imputation and phasing algorithms, based on a comparison of IQS across all combinations (Fig. 10D). There was also a clear relationship between chip density and imputation accuracy, as measured by concordance; as chip density increased, imputation accuracy improved. Omni chip had the greatest chip density and accuracy and the customized chip the lowest (Figs. 10C, 11). From the shape of the chip distributions, we see that the vast majority of the Omni dataset was imputed with very high concordance, whereas less of the Affymetrix input dataset and much less of the Customized chip dataset was imputed with similar accuracy. We also compared imputation accuracy by sex as a check to ensure our QC process does not introduce any artificial differences. Sex had no effect on imputation accuracy for autosomal chromosome 20 (Fig. 10A). Accuracy for females was on average 0.9907 ± 0.0078 while for males it was 0.9906 ± 0.0080.

**Fig. 11.**
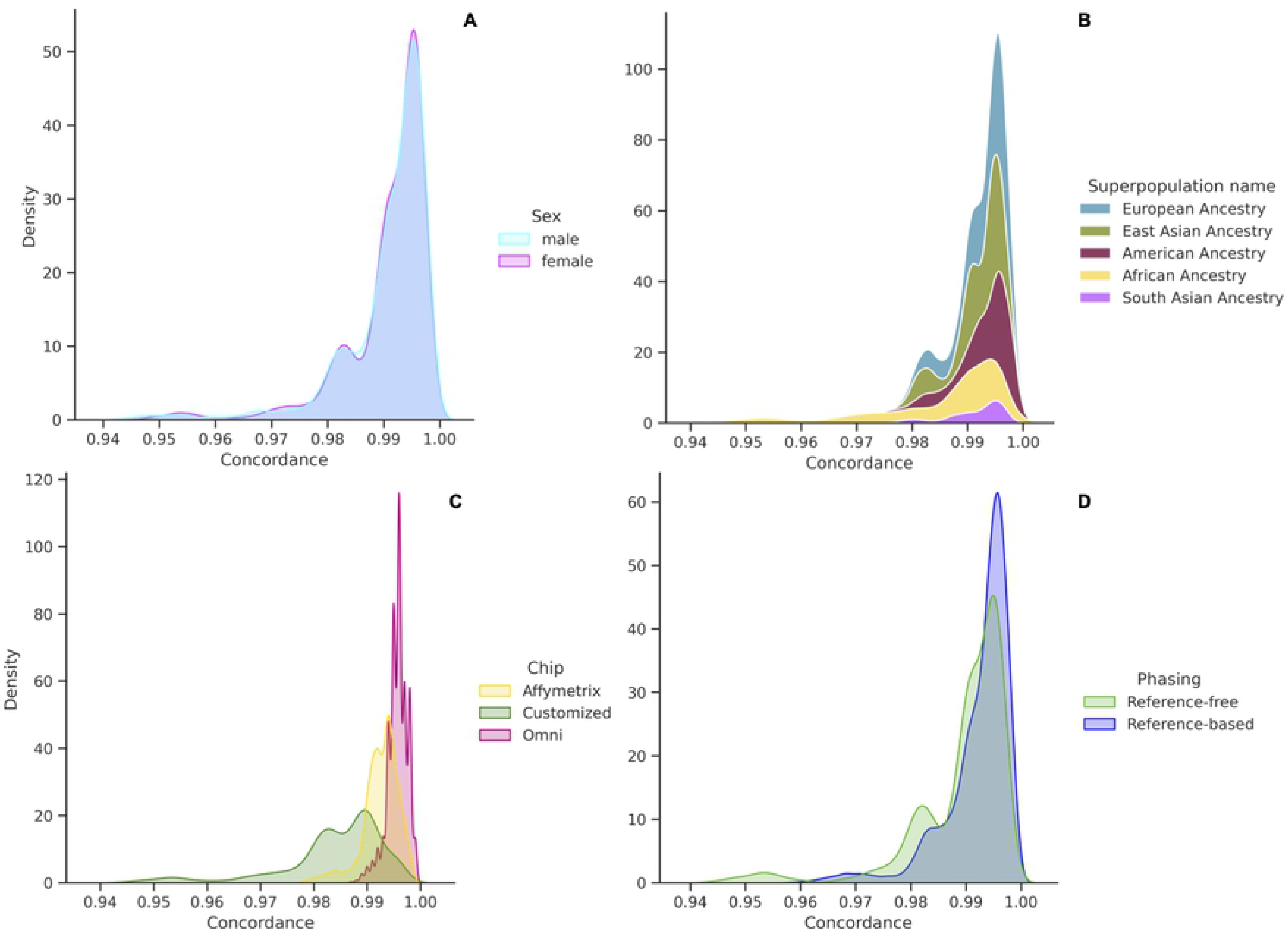
Clustermap of target population against 54 software-reference panel-dataset combinations. This figure depicts the concordance results for the reference-free and reference-based phasing approaches for each of these combinations. Higher density chips with a reference-based phasing approach and with populations without African ancestry obtained better results in terms of imputation accuracy measured by IQS.

### Speed and Memory usage

Of the imputation software’s, Minimac4 appeared to be the most computationally efficient in terms of memory but had the slowest run time, followed by Beagle5.2 and Impute5 (Fig. 12B). Memory usage for Impute5 increased drastically with the size of the input dataset used, while Beagle and Minimac4 were not significantly affected (Fig. 12D). Beagle5.2 had the shortest run time, followed by Impute5 and Minimac4 (Fig. 12A). During phasing, Eagle2.4.1 and ShapeIT4 used less memory than Beagle5.2 and were less affected by the input size of the chip (Fig. 12C). Averaged across the datasets, Eagle2.4.1 was the slowest phasing software while ShapeIT4 was the fastest. Figure 13 shows the average computational run time for each combination. Phasing with ShapeIT4 and imputing with Beagle5.2 was the fastest combination, while phasing with Eagle2.4.1 and imputing with Minimac4 was the slowest.

**Fig. 12.**
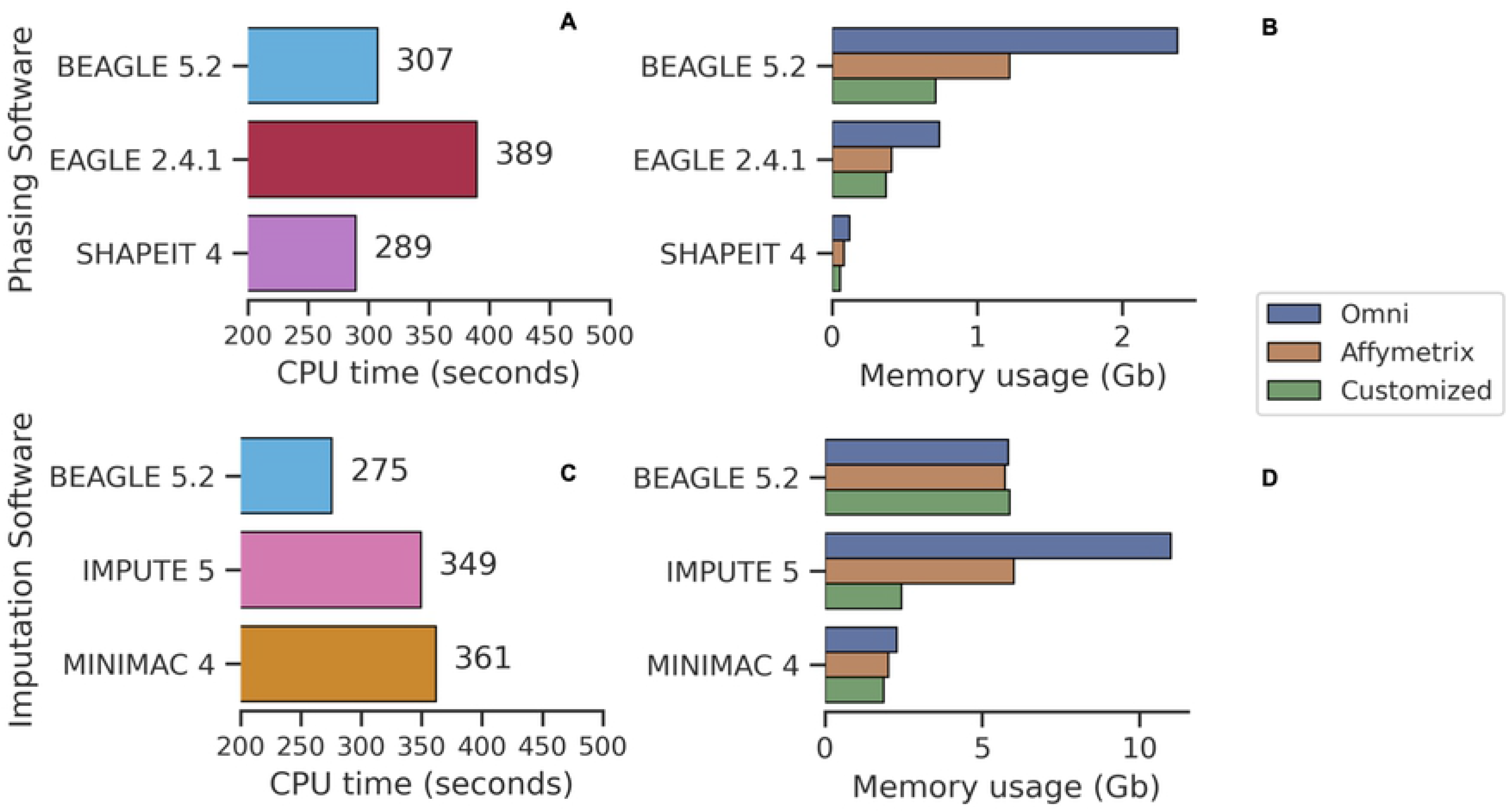
CPU run time and memory usage of imputation and phasing softwares. Average run time for phasing (A) and imputation (C) tools. Average memory usage for phasing (B) and imputation (D) tools.

**Fig. 13.**
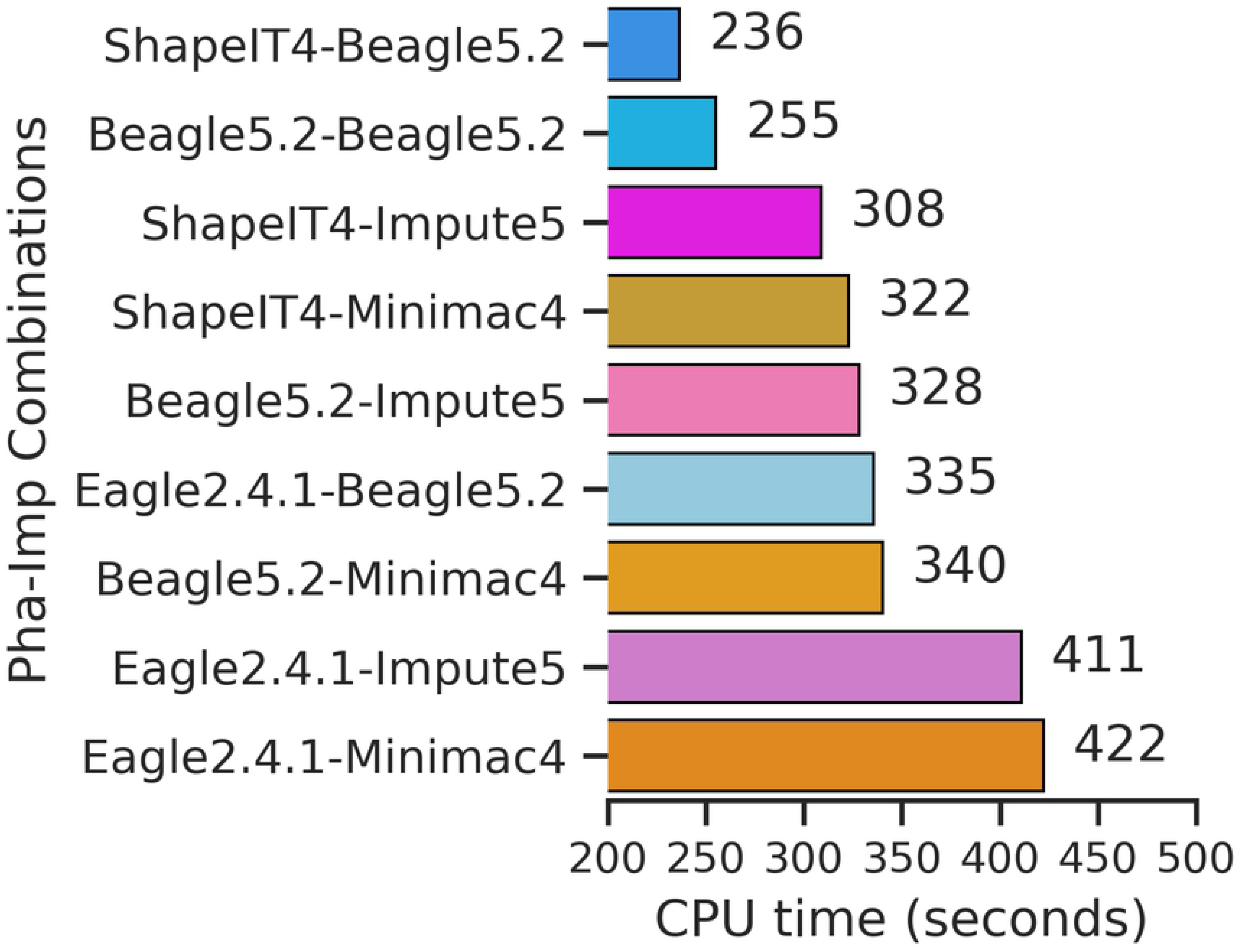
CPU run time of imputation and phasing combinations tested. Average run time for each of the 9 phasing and imputation software combinations.

## Discussion

We performed a rigorous comparison of the most popular phasing and imputation tools currently used by genomics research groups to examine how the process of genotype imputation is affected by different factors, including the choice of reference panel, population, chip density, and allele frequency, with the factor of sex as a control on our process. We also compared the computational load of these different tools and software combinations.

### Factors Affecting Imputation Accuracy

Imputation accuracy decreased with chip density; the Affymetrix chip resulted in lower accuracy than the Omni chip and the customized chip had the lowest imputation accuracy. While this was expected, it also shows how our processing and comparison pipeline may help researchers design better chips by choosing the number and distribution of SNPs for each specific population, and assessing the impact of density and SNP choice on phasing and imputation accuracy; it can also be used to determine whether different sets of chips are likely to perform better with certain combinations of phasing and imputation tools.

Next, we assessed both reference-free and reference-based phasing. Although reference-free phasing was less accurate, we found that increasing chip density alleviates the degree of effect that the lack of reference has on phasing. The difference between reference-free and reference-based phasing was not extreme, suggesting that reference-free phasing may be acceptable in the absence of a representative reference panel. Previous studies comparing phasing accuracy with and without the use of a reference panel have shown that reference-free phasing, such as with Eagle2, can even lead to higher accuracy in cases where the reference panel ancestry and populations do not match well with that of the sample individuals (Loh et al. 2016).

Furthermore, the choice of the reference panel affects imputation accuracy, across all imputation metrics utilized. We note that using the 30X reference panel results in slightly lower imputation accuracy for uncommon variants; this was due to the panel containing more rare SNPs. As variants with lower MAF are more difficult to impute, and are imputed with greater uncertainty and reduced accuracy, these results are expected.

Accuracy was further affected by population but not by sex. Different populations are characterized by differences in LD as a result of differences in genealogical history, and thus have different characteristic LD blocks and LD block sizes, which affect imputation accuracy (David M. Evans 2005). We expect that lower imputation accuracy seen in individuals of AFR ancestry is attributable to the smaller LD blocks characteristic of AFR ancestry, which make it more difficult to correctly impute genotypes.

In agreement with previous research (Shi et al. 2018), we found that variants with low allele frequency are generally imputed poorly. In general, imputation works poorly for variants with low MAF as a function of both bias in the reference panels and bias in the software (Shi et al. 2018). We can address reference-associated bias by significantly increasing the size of the chosen reference panel and including sufficient population-specific samples in the reference. However, addressing software bias would require developing improved imputation algorithms.

Finally, the choice of statistics is important when examining the imputation accuracy of rare and low frequency variants. We found that IQS and squared correlation produced similar means and standard deviations, though this does not necessarily represent similarity of values for particular SNPs. For rare and low frequency variants, concordance rates produce inflated assessments of accuracy (Lin et al. 2010). The higher concordance rate values could mislead a researcher into assuming that these variants were imputed well. However, accuracy for less common variants is best measured using IQS and R^2^ (Ramnarine et al. 2015).

### Choice of Phasing and Imputation Tools

There was a discrepancy in accuracy based on different metrics. Highest average concordance rate was achieved by Beagle5.2 at 0.986, followed by Impute5 and Minimac4, using a reference-based approach during phasing, with the highest density chip dataset as input. In general, choosing Beagle5.2 for imputation and ShapeIT4 for phasing tends to get highly accurate results and is computationally faster. When looking to improve the imputation of rare variants, however, researchers may want to use a mix of Beagle5.2 and Impute5 by applying Beagle5.2 to common variants and Impute5 to rare ones. Impute5 tends to perform better on rare variants, because unlike Beagle5.2, which computes clusters of haplotypes and does its calculations based on them, Impute5 searches the whole space of haplotypes. This is more effective when imputing uncommon variants, but there is a tradeoff of increased computational load.

On the other hand, we see imputation accuracy for Beagle5.2 is better than Impute5 for the filtered phase3 reference panel; this is also expected since the phase 3 panel has fewer rare alleles. Beagle5.2 was also the most stable tool to use across different input sizes. Minimac4 requires the least amount of memory but takes the longest time, which can be a good tradeoff depending on the purpose of the imputation. If the memory usage is limited, and the loss of accuracy is acceptable, then Minimac4 may be the optimal choice of imputation software. It is also important to note that the default parameters have been used for all software. For example, we could reduce the computational load of Impute5 by using parallel processing but this can negatively affect the accuracy results; this negative impact is sufficient to reduce Impute5’s accuracy to below that of Beagle5.2. In conclusion, Beagle might have the best tradeoff between imputation quality and computational efficiency.

In conclusion, differences in imputation and phasing performance may be useful in determining the choice of imputation and phasing tool, depending on the intended downstream usage of the imputed results. However, this study also highlights that current tools are not accurate enough to impute rare and ultra-rare variants, showing that, when corrected for chance concordance and MAF bias, they result in only acceptable imputation accuracy and that there is significant scope for improvement.

## Supporting Information

**S1 Table.**
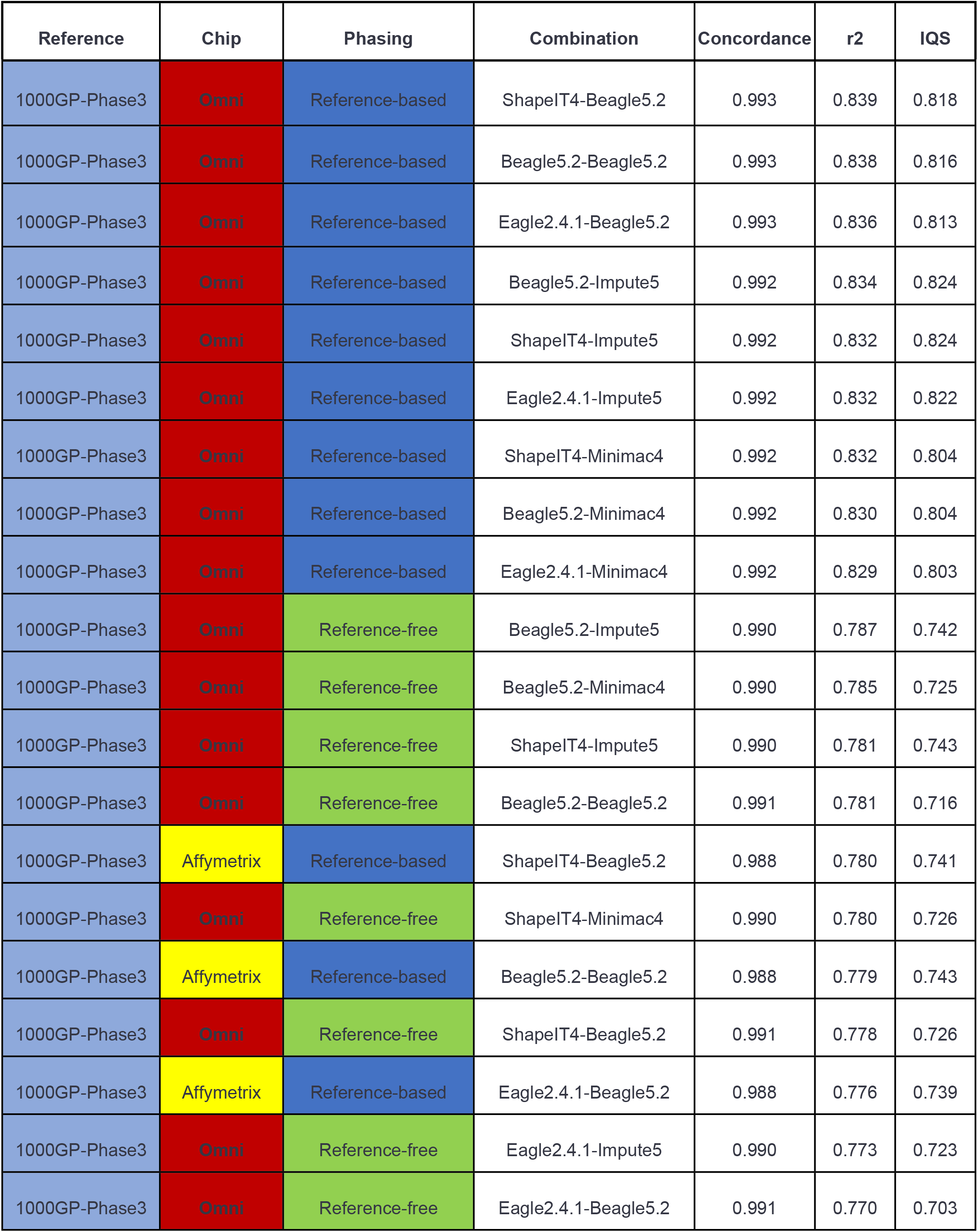
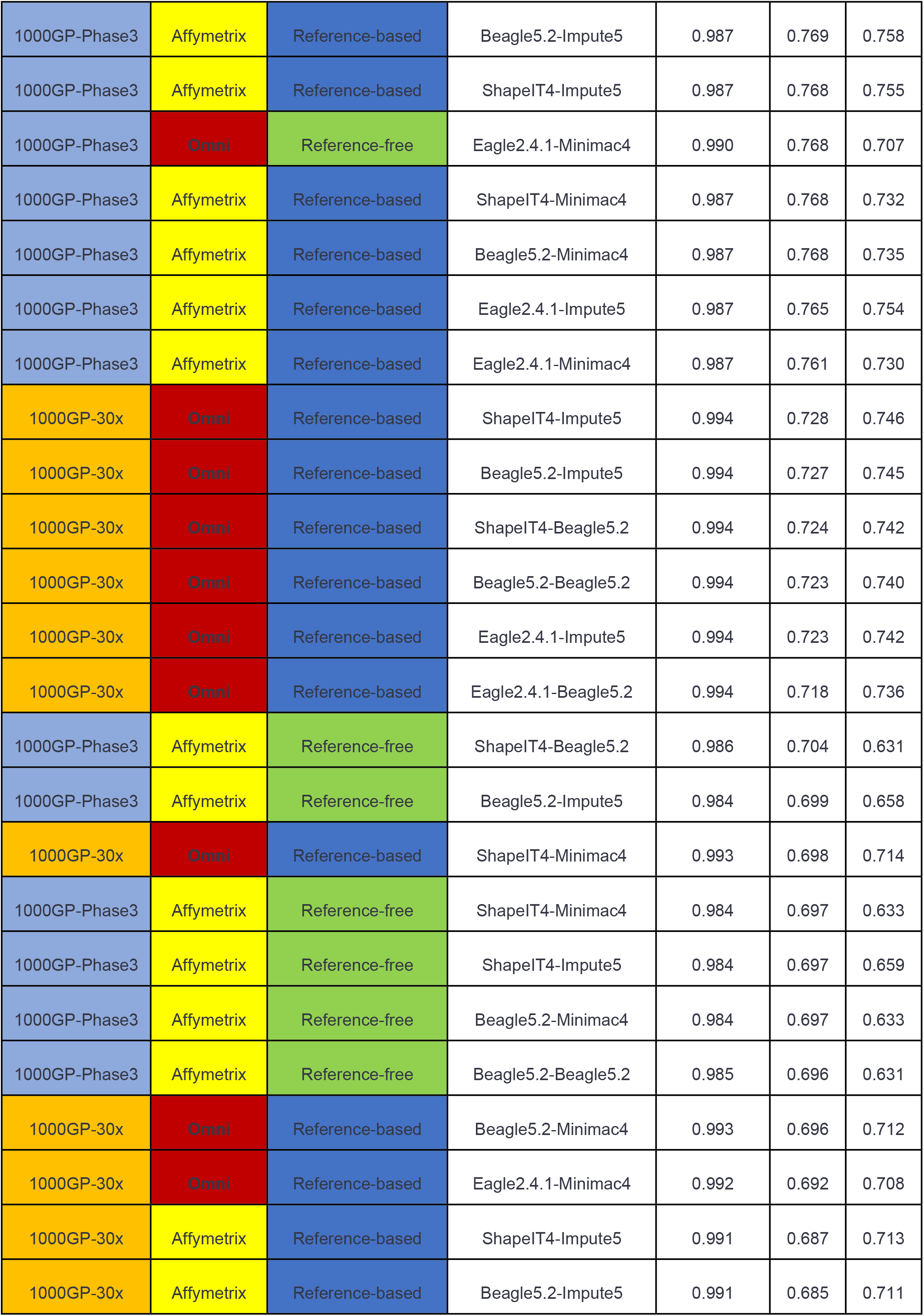
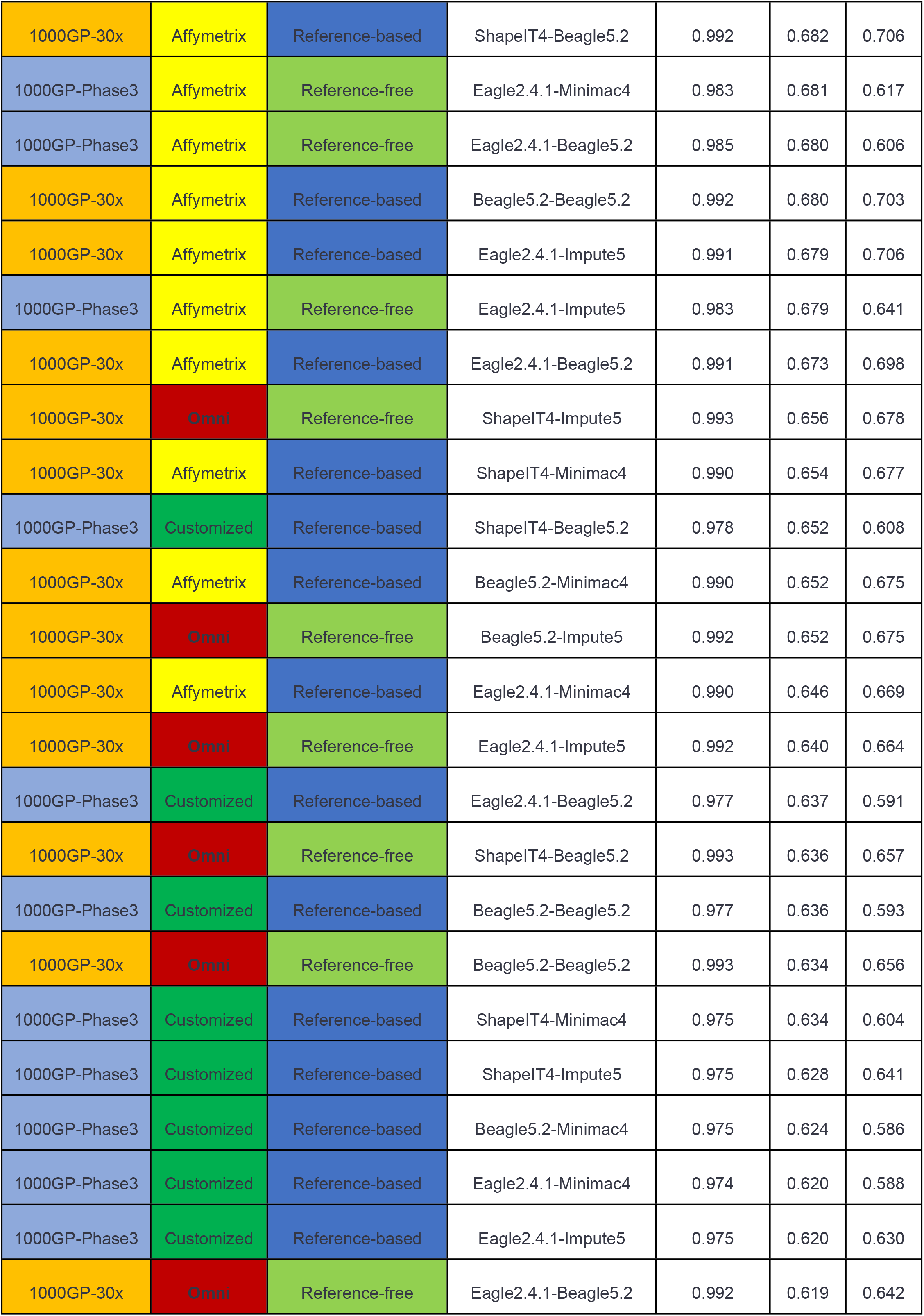
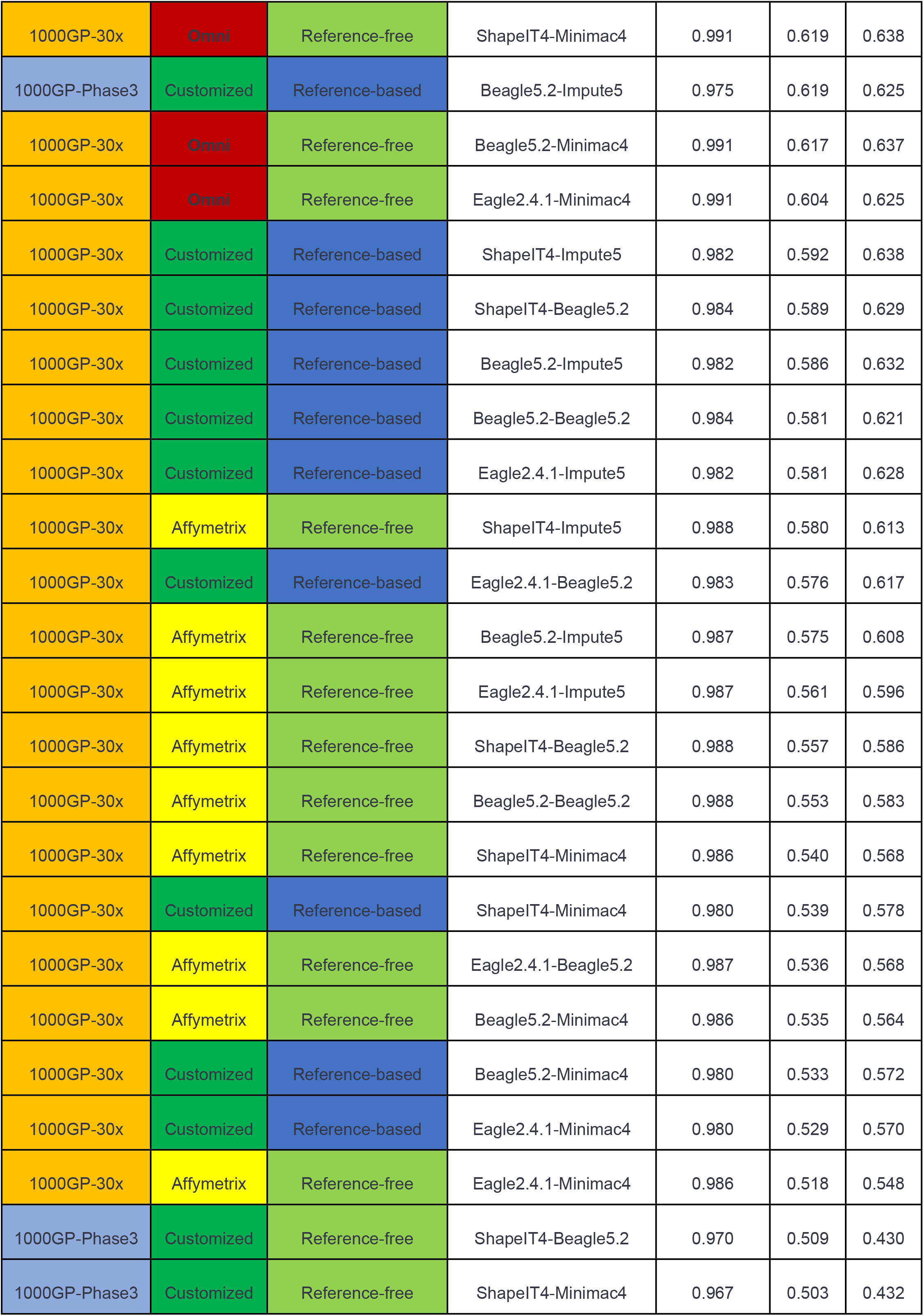
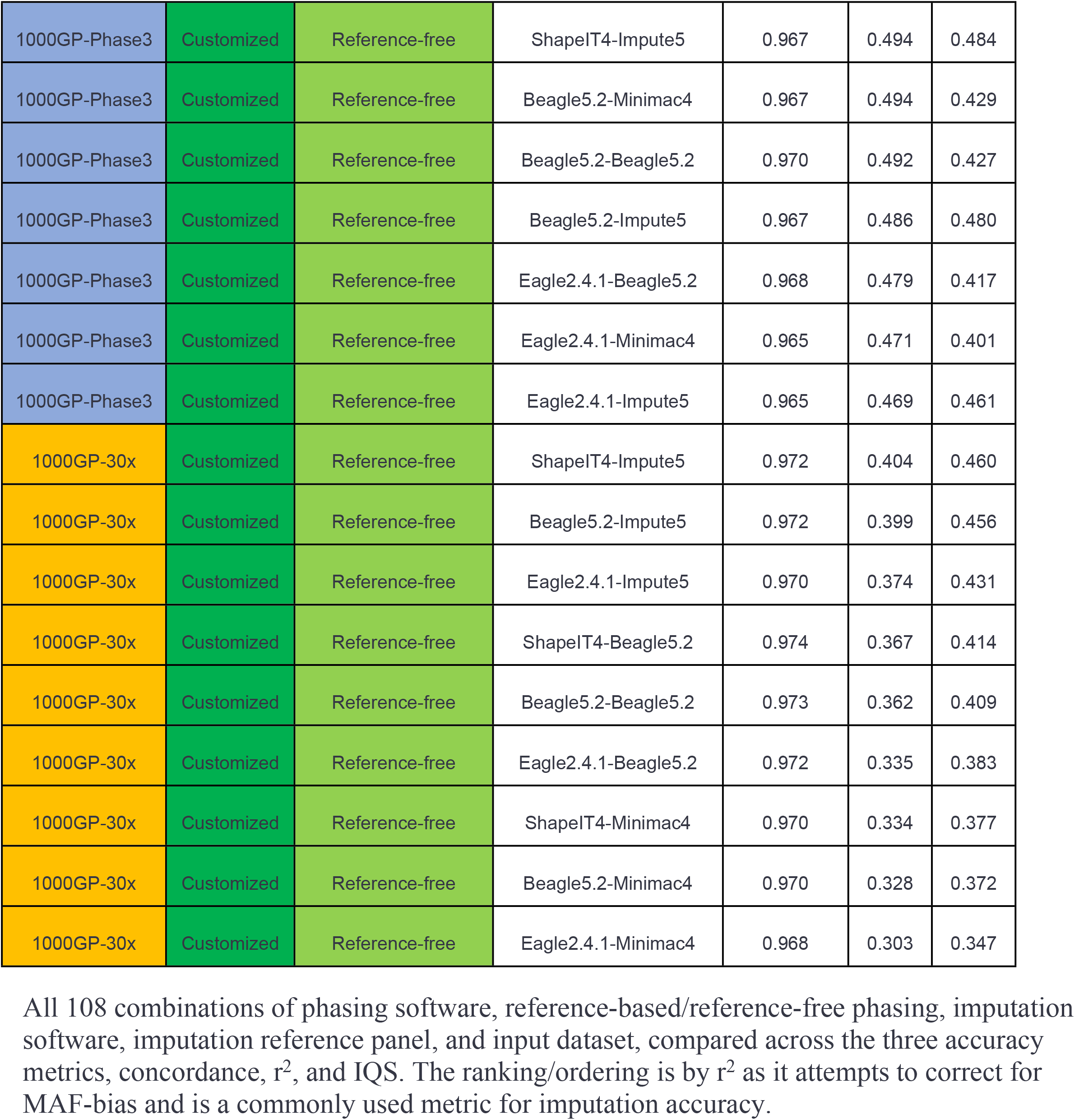
Comparison of all combinations of phasing and imputation tool, reference panel, phasing approach, and chip dataset.

